# Oxidative stress drives potent bactericidal activity of pyrazinamide against *Mycobacterium tuberculosis*

**DOI:** 10.1101/2024.12.17.628853

**Authors:** Nicholas A. Dillon, Elise A. Lamont, Muzafar A. Rather, Anthony D. Baughn

## Abstract

Pyrazinamide (PZA) is a critical component of tuberculosis first-line therapy due to its ability to kill both growing and non-replicating drug-tolerant populations of *Mycobacterium tuberculosis* within the host. Recent evidence indicates that PZA acts through disruption of coenzyme A synthesis under conditions that promote cellular stress. In contrast to its bactericidal action *in vivo*, PZA shows weak bacteriostatic activity against *M. tuberculosis* in axenic culture. While the basis for this striking difference between *in vivo* and *in vitro* PZA activity has yet to be resolved, recent studies have highlighted an important role for cell-mediated immunity in PZA efficacy. These observations suggest that host-derived antimicrobial activity may contribute to the bactericidal action of PZA within the host environment. In this study we show that the active form of PZA, pyrazinoic acid (POA), synergizes with the bactericidal activity of host-derived reactive oxygen species (ROS). We determined that POA can promote increased cellular oxidative damage and enhanced killing of *M. tuberculosis*. Further, we find that the thiol oxidant diamide is also able to potentiate PZA activity, implicating thiol oxidation as a key driver of PZA susceptibility. Using a macrophage infection model, we demonstrate the essentiality of interferon-γ induced ROS production for PZA mediated clearance of *M. tuberculosis*. Based on these observations, we propose that the *in vivo* sterilizing activity of PZA can be mediated through its synergistic interaction with the host oxidative burst leading to collateral disruption of coenzyme A metabolism. These findings will enable discovery efforts to identify novel host- and microbe-directed approaches to bolster PZA efficacy.

## Introduction

Pyrazinamide (PZA) is a cornerstone drug of the first-line therapy for tuberculosis (TB). Following its initial discovery as an antitubercular agent, PZA was administered as monotherapy in humans due to the paucity of effective treatment options (Yeager, Munroe et al. 1952). As a monotherapy, PZA treatment led to initial uniform improvement of clinical symptoms with approximately one-third of patients showing no detectable tubercle bacilli in their sputum (Yeager, Munroe et al. 1952). Subsequent clinical trials involving combination therapy with rifampicin, isoniazid and ethionamide demonstrated that inclusion of PZA could dramatically reduce treatment duration and disease relapse rates (British-Medical-Research-Council 1974, Ormerod and Horsfield 1987). These benefits have been attributed to the unique ability of PZA to kill non-replicating, drug-tolerant populations of bacilli (Hu, Coates et al. 2006) as well as its ability to penetrate and accumulate within granulomatous lesions (Prideaux, Via et al. 2015) and within the phagosomes of infected macrophages (Santucci, Greenwood et al. 2021, Santucci, Aylan et al. 2022). Based on its unique role in current TB therapy, PZA is anticipated to be a component of future regimens for treatment of drug-susceptible and drug-resistant TB infections (Tweed, Dawson et al. 2019).

Despite the fundamental role of PZA in TB therapy, the mechanisms that govern its antitubercular activity are not fully understood. It is known that PZA is a prodrug that must be converted to pyrazinoic acid (POA) by the *Mycobacterium tuberculosis* nicotinamidase PncA (Sreevatsan, Pan et al. 1997). Since PncA activity is non-essential for *in vivo* fitness of *M. tuberculosis* (Boshoff, Xu et al. 2008, Vilchèze, Weinrick et al. 2010), the principal mechanism for clinical PZA resistance is through loss-of-function mutations in *pncA* (Sreevatsan, Pan et al. 1997). Unlike other antitubercular agents, PZA shows a unique conditional bacteriostatic activity *in vitro* against *M. tuberculosis* under environments that promote cell envelope stress, such as low pH (Thiede, Dillon et al. 2022). In contrast to its bacteriostatic action in laboratory culture, PZA shows bactericidal activity in murine models of infection involving immune competent mice (Lanoix, Lenaerts et al. 2015). Further, removal of anti-inflammatory signals through blockade or deletion of IL-10 has been shown to enhance PZA-mediated clearance of *M. tuberculosis* (Dwivedi, Gautam et al. 2023). In contrast, PZA bactericidal activity is abolished in models of TB infection in which cell-mediated immunity has been impaired or dysregulated (Almeida, Tyagi et al. 2014, Lanoix, Lenaerts et al. 2015, Lanoix, Ioerger et al. 2016), highlighting a fundamental connection between PZA action and host response.

Mounting evidence demonstrates that PZA acts through disruption of coenzyme A biosynthesis (Lamont, Dillon et al. 2020). Biochemical and biophysical studies indicate that coenzyme A disruption occurs via interaction of POA with aspartate decarboxylase, PanD (Gopal, Nartey et al. 2017, Gopal, Sarathy et al. 2020, Sun, Li et al. 2020). Consequently, supplementation of *M. tuberculosis* with coenzyme A intermediates can antagonize PZA action and mutations within the *panD* gene have been associated with resistance to POA and PZA (Dillon, Peterson et al. 2014, Shi, Chen et al. 2014, Gopal, Nartey et al. 2017, Gopal, Tasneen et al. 2017, Ramirez-Busby, Rodwell et al. 2017). Despite these key advancements in our understanding of the antitubercular activity of PZA, how these findings relate to the unique bactericidal action of this drug within the host environment has yet to be resolved.

As an intracellular pathogen, *M. tuberculosis* exploits host immune defenses to establish a niche environment for its survival and propagation. Upon inhalation of infectious aerosols by the host, *M. tuberculosis* is deposited within alveolar sacs and is engulfed by alveolar macrophages. Through secretion of effector molecules, *M. tuberculosis* arrests phagosomal maturation and prevents phagosome-lysosome fusion (Pethe, Swenson et al. 2004). Modulation of this initial intracellular environment permits continued bacterial growth through impairment of phagosomal acidification and limited production of reactive oxygen species. In a healthy host, the onset of adaptive immunity marks the end of the acute phase of infection. Through predominantly T_h_1 inflammatory responses driven by interferon-γ (Nathan, Murray et al. 1983, Flynn, Chan et al. 1993) and tumor-necrosis factor-α (Chan, Xing et al. 1992, Flynn, Goldstein et al. 1995), activated macrophages undergo phagosomal maturation and induce phagosome lysosome fusion. Once activated, macrophages increase their production of reactive oxygen and nitrogen species resulting in abatement of bacterial growth and containment of the bacilli within granulomatous lesions.

Production of reactive oxygen species (ROS) is a critical mechanism for control of *M. tuberculosis* within human phagocytes. Superoxide is generated within the phagosome through the transfer of electrons from NADPH to molecular oxygen mediated by the multi-protein complex NADPH phagocyte oxidase. Inside the phagosomal environment, superoxide dismutates to the more stable hydrogen peroxide (H_2_O_2_), which can react with metal ions to produce highly reactive hydroxyl radicals (^•^OH). ^•^OH induces microbial death through broad oxidative damage to proteins, DNA, and lipids. Defects in the ROS producing NADPH phagocyte oxidase complex are associated with increased susceptibility to mycobacteria in humans (Bustamante, Aksu et al. 2007, Lee, Chan et al. 2008).

Prior studies in cell-free systems have established that H_2_O_2_-mediated oxidation of hydrocarbons can be accelerated in the presence of POA-iron complexes (Shulpin, Attanasio et al. 1993). While the chemical mechanism governing this oxidation is unclear, it is predicted to involve the direct formation of ^•^OH from the interaction of H_2_O_2_ with POA-coordinated iron (Kirillov and Shul’pin 2013). Whether POA-mediated formation of ^•^OH is associated with the *in vivo* bactericidal action of PZA has not been reported. However, Lien and colleagues recently demonstrated that deletion of a novel mycobacterial nanocompartment involved in oxidative stress defense sensitized *M. tuberculosis* to PZA in a murine model of infection (Lien, Dinshaw et al. 2021). In this report, we investigate the association between PZA susceptibility and oxidative stress in *M. tuberculosis* to delineate the role of the host environment in bactericidal action. We demonstrate that normally sub-inhibitory concentrations of PZA and H_2_O_2_ can act synergistically against *M. tuberculosis* resulting in bactericidal activity. Moreover, we demonstrate that potentiation of PZA action is driven through thiol stress. Finally, we find that this activity correlates with the bactericidal action of PZA within infected macrophages. Taken together, these observations support a role for host-induced oxidative stress in the *in vivo* bactericidal action of PZA.

## Results

### Pyrazinamide and hydrogen peroxide show synergistic bactericidal activity against *M. tuberculosis*

Given the critical role of host-derived ROS in controlling tuberculosis infections and the diminished antitubercular efficacy of PZA in infection models with impaired macrophage activation (Almeida, Tyagi et al. 2014, Lanoix, Lenaerts et al. 2015, Lanoix, Ioerger et al. 2016), we explored the potential interaction between PZA and ROS on *M. tuberculosis* in culture. *M. tuberculosis* was grown in 7H9 medium (pH 5.8) supplemented with 10% ADS, 0.2% glycerol and 0.05% tyloxapol. At mid-exponential phase, PZA (200 µg/mL) or vehicle (DMSO) was added and cultures were incubated for an additional 3 days. Aliquots of cultures were then exposed to a geometric concentration series of ROS generating compounds (H_2_O_2_, menadione or clofazimine) for 24 hrs. Cells were serially diluted and plated on 7H10 agar to determine the titer of surviving colony forming units (CFU). PZA pretreated cultures were significantly more susceptible to H_2_O_2_ mediated killing with a ≥16-fold decrease in the minimum bactericidal concentration (MBC, minimum concentration required to kill >99% of bacterial cells) relative to the no PZA control (Figure 1A). This combinatorial approach was also applied with menadione and clofazimine which have been reported to interact with the mycobacterial respiratory chain resulting in superoxide production (Akhtar, Srivastava et al. 2006, Yano, Kassovska-Bratinova et al. 2011). While both menadione and clofazimine were also found to elicit bactericidal activity against *M. tuberculosis* as previously reported, neither showed the same level of enhanced activity against PZA-treated cells (Figure 1B, C).

**Figure 1.**
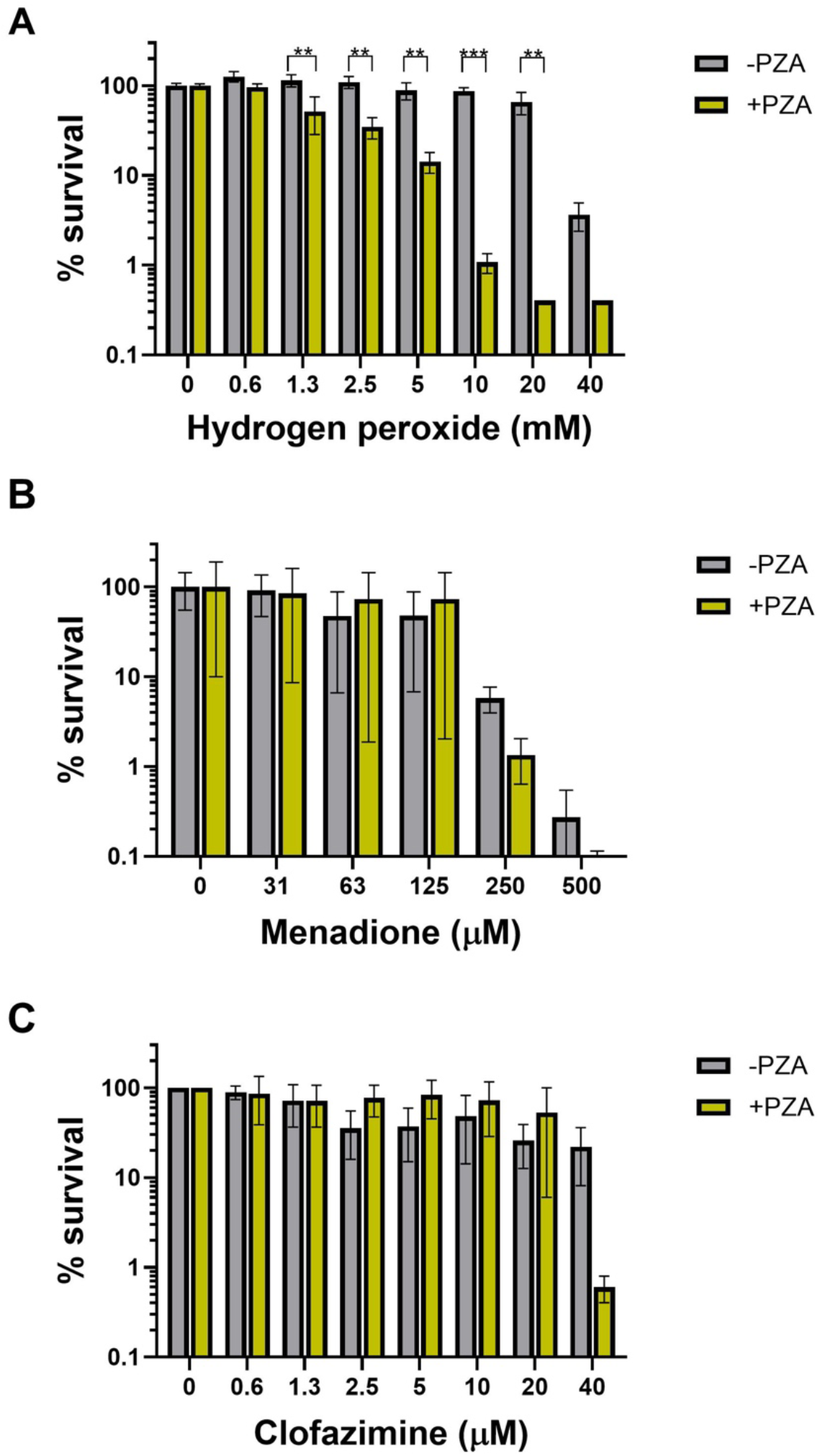
PZA treatment synergizes with H_2_O_2_ to kill *M. tuberculosis*. Bactericidal activity of ROS generators was examined after exposure to 200 µg/mL PZA for (A) H_2_O_2_, (B) clofazimine and (C) menadione. Data shown represent the mean and standard deviation from biological triplicates. Statistical significance was calculated with a multiple comparisons two-way ANOVA, \**P*≤0.05, \*\**P*≤0.01.

To further examine the extent of interaction between H_2_O_2_ and PZA, checkerboard assays (Odds 2003) were conducted with *M. tuberculosis* to determine the fractional inhibitory concentration index (FICI) values for this combination. Similar to the prior experiment, cells were exposed to PZA in 7H9 for 72 hours prior to H_2_O_2_ exposure. Cultures were then incubated for 7 days and growth was assessed by measuring OD_600_. Cultures that showed less than 10% growth relative to the no drug control were considered inhibited. The interaction of PZA and H_2_O_2_ was found to be strongly synergistic (FICI ≤ 0.5) against strains H37Rv (FICI of 0.19) and H37Ra (FICI of 0.06) (Figure 2A, 2B).

**Figure 2.**
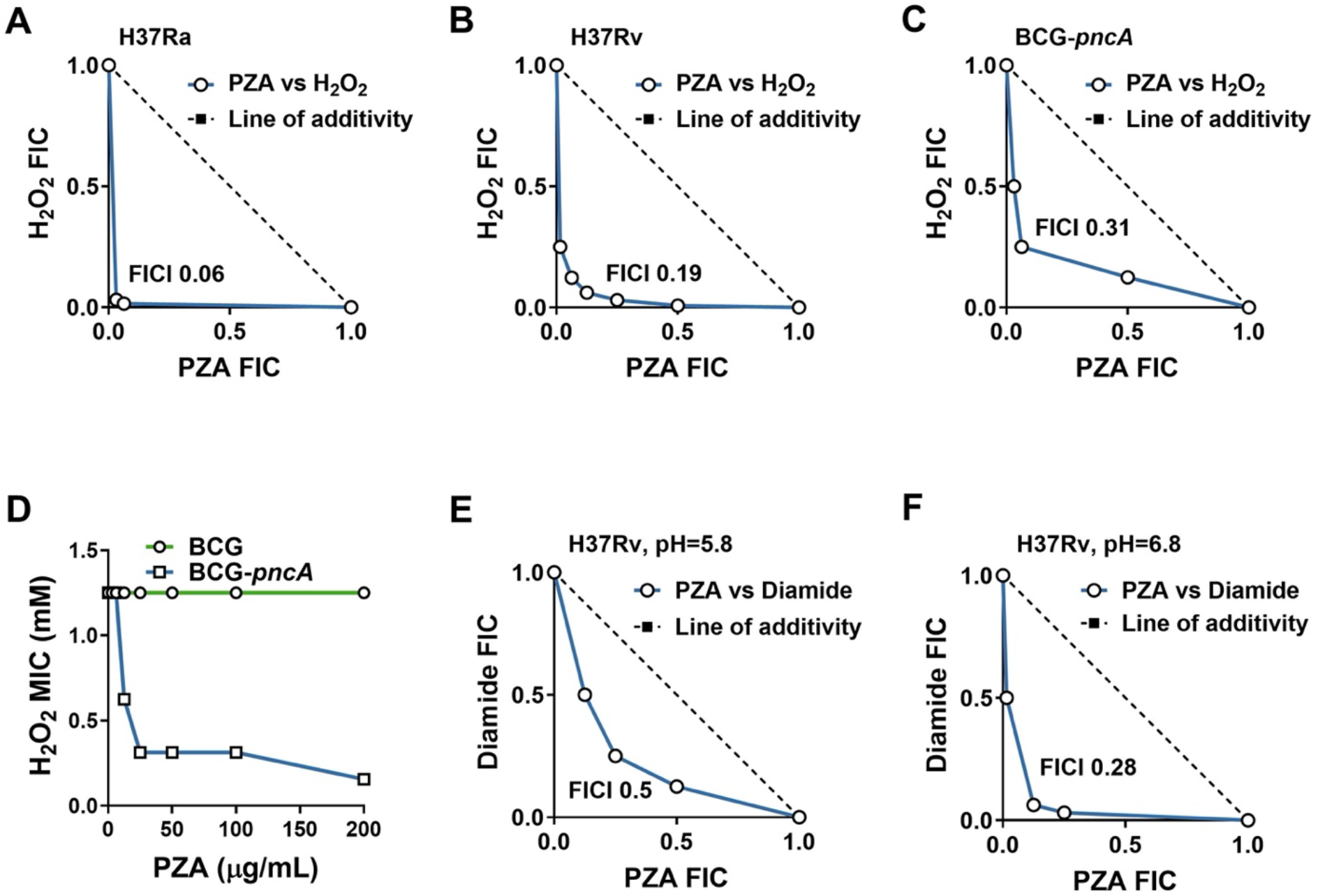
Antimycobacterial action of PZA is potentiated by oxidative stress. Synergy between PZA and oxidizing agents H_2_O_2_ and diamide was assessed against *M. tuberculosis* H37Ra (A), *M. tuberculosis* H37Rv (B, E, F), and *M. bovis* BCG with and without a functional copy of *pncA* (C, D). Cultures were exposed to PZA for 72 hours prior to treatment with oxidizing agents. FIC plots were generated following outgrowth from challenge with H_2_O_2_ (A-D) or diamide (E, F) in 7H9 with ADS pH 5.8 (A-E) or 6.8 (F). Lowest measured FICI values are show within each plot. Data represent the mean of three biological replicates.

Since the antitubercular action of PZA is dependent upon PncA-mediated conversion to POA, we sought to determine whether PncA activity was essential for the observed synergy with H_2_O_2_. *Mycobacterium bovis* is naturally resistant to PZA due to a loss-of-function mutation in the *pncA* gene rendering this species incapable of generating POA (Sreevatsan, Pan et al. 1997). Ectopic expression of a functional copy of *pncA* enables POA conversion thereby restoring PZA susceptibility in the attenuated *M. bovis* vaccine strain Bacillus Calmette Guérin (BCG) Pasteur (Baughn, Deng et al. 2010). Checkerboard assays with PZA and H_2_O_2_ were carried out in *M. bovis* BCG and BCG-*pncA* to assess the role of POA conversion in synergy between PZA and H_2_O_2_. Like previous observations for *M. tuberculosis*, PZA was synergistic with H_2_O_2_ against the BCG-*pncA* strain (FICI of 0.31) (Figure 2C). However, synergy with H_2_O_2_ was not observed for the parental BCG strain, indicating that conversion of PZA to POA is essential for synergy between H_2_O_2_ and PZA (Figure 2D).

Since oxidative stress results in broad oxidation of cellular thiols within biological systems (Baba and Bhatnagar 2018) and PZA is known to disrupt synthesis of the thiol cofactor coenzyme A, we were curious to determine whether thiol oxidation is sufficient to potentiate susceptibility of *M. tuberculosis* to PZA. It has previously been shown that diamide can elicit growth-inhibitory thiol stress in *M. tuberculosis*, and this effect is enhanced under conditions of low pH (Coulson, Johnson et al. 2017). Consistent with a role for thiol stress in potentiation of PZA action, co-exposure of *M. tuberculosis* to diamide and PZA in checkerboard format revealed synergy under acidic (pH 5.8, FICI of 0.5) (Figure 2E) and more so under circumneutral (pH 6.8, FICI of 0.28) (Figure 2F) conditions. These observations suggest that thiol oxidation is likely to be a key contributor to the enhanced activity of PZA under conditions of oxidative stress.

### PZA and H_2_O_2_ result in synergistic oxidation of cellular targets

To assess the ability of POA to accelerate intrabacterial release of ^•^OH, we utilized the fluorogenic probe CellRox Green to quantify the abundance of oxidative radicals (Yang and Choi 2018) in *M. tuberculosis* following exposure to PZA and H_2_O_2_. *M. tuberculosis* H37Rv was pretreated with PZA (100 and 200 µg/mL) or vehicle control (DMSO) for 3 days. Cultures were subsequently treated with H_2_O_2_ (0.5-10 mM) for up to 24 hrs. In the absence of H_2_O_2_ there was no discernable increase in oxidative radical production due to the administration of PZA (Figure 3A). However, pretreatment with PZA followed by exposure to H_2_O_2_ resulted in a time- and dose-dependent increase in signal relative to the vehicle control (Figure 3B-4D). Together these data suggest that H_2_O_2_ synergizes with PZA through enhanced intrabacterial release of ^•^OH.

**Figure 3.**
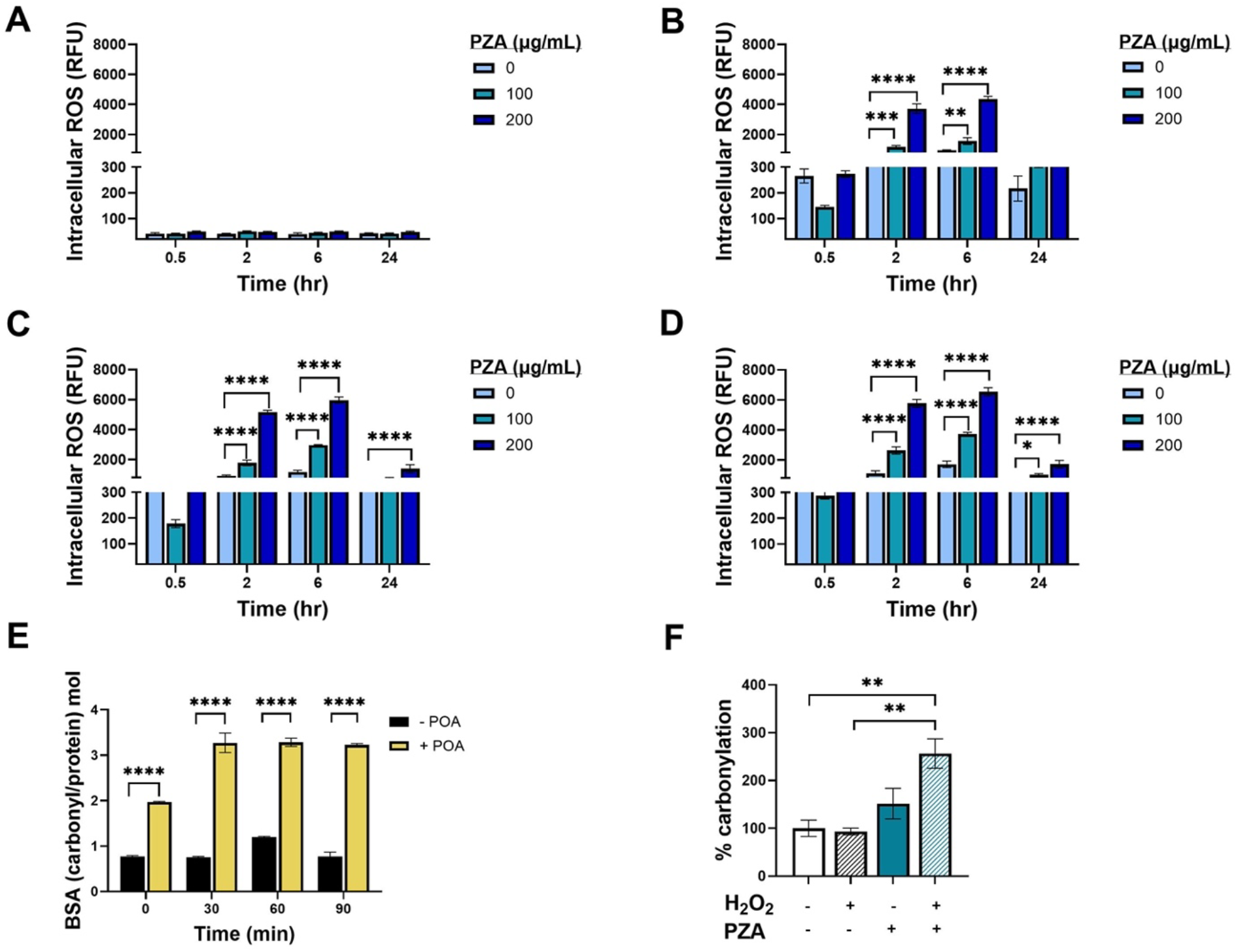
PZA treatment enhances H_2_O_2_ associated intracellular ROS production and oxidative damage in *M. tuberculosis*. Intracellular ROS levels were measured by CellRox Green assay using mid-exponential phase *M. tuberculosis* H37Rv cultures which had been treated with either no PZA, 100 µg/mL PZA, or 200 µg/mL PZA for 3d at pH 6.8 prior to the addition of (A) no H_2_O_2_, (B) 0.5 mM H_2_O_2_, (C) 5 mM H_2_O_2_, or (D) 5 mM H_2_O_2_. Protein carbonylation was examined as a biological marker of oxidative damage. Carbonyl groups were derivatized with 2,4-dinitrophenylhydrazine and the molar ratio of carbonyl groups to protein was determined for (E) 200 µg/mL of BSA with 1 mM Fe^2+^, 25 mM H_2_O_2_ in 10 mM MES buffer (pH 5) with either no POA or 2 mM POA, or (F) mid-exponential phase *M. tuberculosis* H37Ra cultures which had been treated with either no drug or PZA at 200 µg/mL PZA for 3d at pH 5.8, cells were then subsequently exposed to no H_2_O_2_ or 5 mM H_2_O_2_ for 24 hrs. All experiments were conducted in triplicate. Statistical significance was calculated with a multiple comparisons two-way ANOVA, *≤0.05, \*\**P*≤0.01, ***P≤0.001, ****P≤0.0001.

Through damaging multiple cellular components, host-derived ROS plays a key role in eliminating pathogenic bacteria. If POA activity is mediated through ROS, then PZA treated cultures of *M. tuberculosis* should have increased levels of oxidative damage. Protein carbonylation, which occurs when protein side chains become carbonylated upon exposure to ROS, is a biologically quantifiable marker of oxidative damage. Carbonyl groups can be derivatized with 2,4 dinitrophenylhydrazine (DNPH) and then quantified by measuring the adduct abundance in a given protein sample spectrophotometrically. An assay using bovine serum albumin (BSA), Fe^2+^, and H_2_O_2_ was implemented to assess POA associated protein carbonylation in a cell-free system. Within 30 minutes after the addition of H_2_O_2_, BSA associated carbonylation was increased approximately 4-fold by POA compared to baseline (Figure 3E). To measure carbonylation from whole cells, *M tuberculosis* H37Ra was treated with PZA at pH 5.8 for 72 hours, challenged with H_2_O_2_ for 24 hours, cultures lysed, and extracted for total protein that was later derivatized with DNPH. PZA treatment significantly increased protein carbonylation in *M. tuberculosis* when compared to the untreated (2.6-fold, p=0.0077) and H_2_O_2_ only (2.8-fold, p=0.0060) controls (Figure 3F). Interestingly, PZA treatment alone in the absence of H_2_O_2_ trended to higher, yet statistically insignificant, levels of protein carbonylation than either the untreated (1.5-fold, p=0.4681) or H_2_O_2_ only (1.6-fold, p=0.3766) controls (Figure 3F). The addition of H_2_O_2_ to PZA treated cells led to an additional 1.7-fold (p=0.0602) increase in protein carbonylation over that observed with PZA only.

### Bactericidal activity of PZA is dependent upon host-derived ROS within macrophages

Previous *in vivo* studies indicate a dependence on cell-mediated responses for PZA bactericidal activity. Based on our findings, we hypothesize that PZA activity against *M. tuberculosis* in the host is mediated through POA-driven sensitization to the oxidative burst in macrophages. To directly test this hypothesis, we examined the requirement for host derived ROS for PZA activity using virulent H37Rv and the minimal unit of TB infection, the macrophage (VanderVen, Huang et al. 2016). We utilized well established macrophage cell line infection models, a murine RAW 264.1 and PMA differentiated human THP-1 macrophages exposed to virulent *M. tuberculosis* H37Rv. Macrophages were either left resting or activated with IFN-γ prior to infection. *M. tuberculosis* H37Rv infected macrophages were treated daily with PZA (400 ug/mL) or DMSO vehicle control and monitored up to 10 days (Figure 4A,B). In both RAW 264.1 and THP-1 cell lines, IFN-γ activation led to an approximate 4-fold reduction in bacterial CFUs compared to resting macrophages (Figure 4A,B). The addition of PZA to IFN-γ activated macrophages decreased the bacterial burden a further ∼1.5-fold, yet, surprisingly, had no discernable effect on *M. tuberculosis* growth in resting cells (Figure 4A,B). IFN-γ signaling activates enhanced anti-mycobacterial macrophage functions, which includes elevated production of ROS (MacMicking 2014, Kak, Raza et al. 2018). We therefore assessed the role of host produced ROS in PZA action by treating *M. tuberculosis* infected IFN-γ activated macrophages with the well-established ROS scavenger, N-acetyl-L cysteine (NAC). In both macrophage subsets the addition of NAC in the context of PZA treatment resulted in the complete abolishment of drug related bactericidal activity (Figure 4A,B). CFU burden in NAC treated cultures were comparable to those of resting macrophages with and without PZA treatment, demonstrating the importance of host-derived ROS for bactericidal activity (Figure 4A,B).

**Figure 4.**
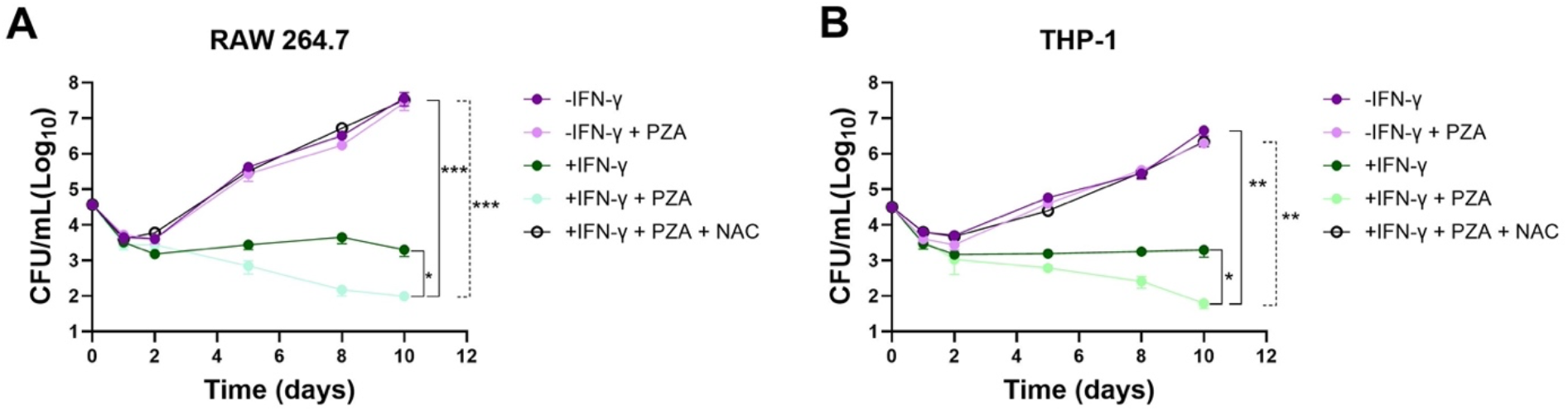
PZA activity is dependent on the production of host derived ROS. PZA bactericidal activity in *M. tuberculosis* H37Rv infected (A) murine RAW 264.1 or (B) human THP-1 macrophages. Macrophages were either untreated or activated through preincubation with IFN-γ, and/or N-acetyl-L-cysteine (NAC) prior to the infection. After 2 hr of infection cells were washed, and PZA was added as indicated. Cultures were plated for CFU enumeration at the indicated timepoints. Statistical significance was calculated with a multiple comparisons two-way ANOVA, *≤0.05, \*\**P*≤0.01, ***P≤0.001.

## Discussion

Despite several decades of PZA use in first-line TB therapy, the underlying basis for its divergent activity against *M. tuberculosis in vitro* and *in vivo* has remained elusive. Recent investigations utilizing murine models of TB infection have suggested that the *in vivo* bactericidal activity of PZA relies on activation of the cell-mediated immune response (Almeida, Tyagi et al. 2014, Lanoix, Lenaerts et al. 2015, Lanoix, Ioerger et al. 2016). These findings indicate that antimicrobial effectors within the host microenvironment are of fundamental importance in the bactericidal action of PZA. Through the use of axenic cultures and macrophage infections, our study demonstrates that ROS plays a pivotal role in driving bactericidal activity of PZA against *M. tuberculosis*. Consistent with the crucial role of pyrazinamidase in PZA activation, we observed that strains with *pncA* loss-of-function become resistant to this bactericidal action. Additionally, we demonstrate PZA treatment leads to increased intrabacterial oxidative damage. The role of ROS in driving PZA bactericidal activity in the host environment was supported by its indispensability in macrophage infection models and highlights the importance of localization of PZA to the phagosome (Santucci, Greenwood et al. 2021, Santucci, Aylan et al. 2022).

IFN-γ activation is a key signal for phagosomal ROS production in monocytic cells. In this study, IFN-γ activation was found to be crucial for bactericidal activity of PZA against intracellular tubercle bacilli. While IFN-γ activation induces various antimicrobial responses in phagocytes, the finding that the ROS scavenger NAC abolishes drug activity allows these other factors to be decoupled from PZA action within this niche. However, contribution of other important oxidative antimicrobial factors such as nitric oxide (Chan, Xing et al. 1992) and metabolic aldehydes (Berry, Espich et al. 2023) cannot be excluded and warrant further investigation. In axenic culture, POA may also interact with endogenously-derived ROS. Previous studies have demonstrated that *M. tuberculosis* is more susceptible to perturbations in endogenous ROS production relative to other mycobacteria (Tyagi, Dharmaraja et al. 2015). In line with these findings, our study demonstrates that PZA-treated cells undergo sublethal oxidative damage even without exogenously supplied H_2_O_2_. However, the abundance of endogenous ROS alone is insufficient to drive bactericidal action of PZA. These observations indicate that the varying ROS levels in different environments likely contribute to the disparate action of PZA in laboratory culture versus the host environment which is in line with the role for peroxide stress in PZA susceptibility in murine infection models (Lien, Dinshaw et al. 2021).

Based on the highly reactive nature of ^•^OH radicals, PZA-mediated enhanced susceptibility to H_2_O_2_ is likely to have pleiotropic effects on *M. tuberculosis* resulting in oxidative damage to lipids, nucleic acids and various central metabolites. In this study, we used protein carbonylation as a readout for cellular oxidative damage. While additional investigation is necessary to define the full extent of molecular interactions at play, the finding that PZA action is potentiated by the thiol oxidant diamide suggests thiol stress as a major mediator.

In the ongoing pursuit of improved anti-mycobacterial therapies, this study reveals new opportunities for potentiating PZA action. Given that immune impairment is both associated with progression of TB disease and in animal models of infection is associated with PZA treatment failure, it is of vital importance to identify means by which to drive PZA action independent of host immune status. Such approaches could include microbe-directed approaches that impair mycobacterial oxidative stress defense or disrupt thiol homeostasis, or could include host-directed approaches that bolster production of host-derived antimicrobial products such as reactive oxygen and reactive nitrogen species or other toxic molecules such as reactive aldehydes.

## Materials and methods

### Bacterial strains and growth media

*M tuberculosis* strains H37Rv and H37Ra, and *M. bovis* strains BCG and BCG-*pncA* were grown at 37°C in Middlebrook 7H9 medium (Difco) supplemented with 0.2% (vol/vol) glycerol, 0.05% (vol/vol) tyloxapol, and either 10% (vol/vol) oleic acid-albumin-dextrose-catalase (OADC) or albumin-dextrose-sodium chloride (ADS) as indicated.

### MIC and FIC determinations

Drug susceptibility was determined by measuring their optical densities at 600 nm in comparison to untreated controls. MIC assays were conducted by determining the minimum amount of drug that was required to inhibit 90% of the growth in treated cultures when compared to untreated controls. POA stocks were made in water and the pH was adjusted with NaOH to match the pH of the medium prior to its addition. Synergies between different combinations of antimicrobials were determined by conducting fractional inhibitory concentration (FIC) experiments. The FIC experiments were carried out by setting up checkerboard assays with varying 2-fold concentrations of each antimycobacterial compound examined. Inhibition was scored in the FIC experiments based on the lowest amount of antimicrobial that was required to inhibit 90% of the growth in untreated cultures during concurrent exposure to a constant concentration of the other antimicrobial being tested. The fractional inhibitory concentration index (FICI) was determined by calculating the sum of the measured FIC values for each compound at each point and then reporting the lowest value determined. All MIC and FIC assays were conducted in 7H9 complete medium with ADS (pH 5.8) at 37°C on a 100 rpm rotary shaker in triplicate.

### Bacterial survival assays

H37Ra cultures grown in complete MB7H9 supplemented with ADS (pH 5.8) were pretreated with PZA (200 µg/mL) for 72 hrs prior to the addition of the indicated concentrations of H_2_O_2_, clofazimine, or menadione for 24 hrs at 37°C with shaking (100 rpm). After 24 hrs cultures were 10-fold serially diluted and spotted on 7H10 complete medium with 10% OADC and incubated at 37°C until colonies were visible. Each experiment was carried out in triplicate.

### Protein carbonylation assays

Protein carbonylation assays were performed using a cell-free system with BSA (200 µg/mL) or using live *M. tuberculosis* H37Ra. The cell-free assay was conducted in 10 mM MES (pH 5) buffer at 37°C. BSA or H37Ra cells were used as protein sources for this assay. POA and Fe^2+^ were added where indicated at 2 mM and 1 mM, respectfully. Protein was harvested at the indicated timepoints after the addition of 25 mM H_2_O_2_.

*M. tuberculosis* H37Ra cultures were grown in 7H9 complete medium with 10% ADS (pH 6.8) to mid-log phase, washed once with 7H9 complete medium containing dextrose-sodium chloride (pH 5.8), and resuspended in 7H9 complete medium (pH 5.8) prior to the addition of PZA. One hundred twenty-five mL of *M. tuberculosis* H37Ra (OD_600_ = 0.05) was used for each sample. Cultures receiving drug treatment were treated with PZA at 200 µg/mL for 72 hours prior to the addition of 5 mM H_2_O_2_ for 24 hours at 37°C with shaking. Cultures were then pelleted by centrifugation at 13,000 *x g* and resuspended in 2 mL of 7H9 complete medium containing dextrose-sodium chloride (pH 5.8) Cultures were then lysed via bead beating to extract protein. Measurement of carbonylation was carried out via the spectrophotometric method as described (Wehr and Levine 2013). In brief, lysates were first treated with 1% streptomycin sulfate for 30 min on ice followed by a 30 min spin at 13,000 rpm at 4°C to pellet DNA. The supernatants were transferred and treated with 2,4 dinitrophenylhydrazine (DNPH) for 10 min prior to protein precipitation using 10% trichloroacetic acid. Protein pellets were then washed thrice with 1 mL of ethanol-ethyl acetate followed by low-speed pelleting. Protein was resuspended in 800 µL of 6.0 M guanidine HCl, 0.5 M potassium phosphate (pH 2.5). DNPH signal was quantified by measuring absorbance at 395 nm using a Genesysis 20 spectrophotometer.

### PZA activity in macrophages

Unless otherwise stated all cell culture incubations were conducted at 37°C in a humidified chamber containing 5% CO_2_. RAW 264.7 murine macrophage and THP-1 monocytic cell lines were generous gifts by Kaylee Schwertfeger (University of Minnesota). RAW 264.7 cells were maintained in Dulbecco’s Minimal Essential Medium (DMEM) with 10% Fetal Bovine Serum (FBS) and 1% pen/strep solution at 37°C. THP-1 cells were maintained in ATCC formulated RPMI 1640 containing 10% FBS, 1% pen/strep solution, and 0.05 mM 2-mercaptoethanol, herein referred to as RPMI 1640 complete. RAW cells were seeded at a density of 2.0 x 10^4^ cells/well in 24 well plates in culture medium and allowed to adhere overnight. THP-1 cells were seeded separately at the same density as RAW 264.7 cells and differentiated for 3 days using phorbol 12-myristate 13-acetate (PMA; 20 nM). PMA was subsequently removed from THP-1 cells by washing cells thrice in Hank’s Buffered Salt Solution. THP-1 cells were allowed to rest for 24 hrs in RPMI 1640 complete. All cells were washed thrice using 1X Dulbecco’s Phosphate Buffered Saline (D-PBS) and incubated for a further 24 hr in appropriate culture medium without antibiotic. Designated cell plates were activated with mouse IFN-γ (5 ng/mL; Shenandoah Biotechnology Inc.) or human IFN-γ (5 ng/mL; Shenandoah Biotechnology Inc.) overnight. Cells were washed as before and resuspended in culture medium without antibiotics. Fifteen minutes prior to infection cells were incubated with N-acetyl-L-cysteine (NAC) (10 mM) or untreated culture medium. Mid-log phase *M. tuberculosis* H37Rv was allowed to infect macrophages at MOI 1 for 2 hr. Upon completion of infection, macrophages were washed as before and DMEM or RPMI 1640 was replenished with the following treatments: control, vehicle control (DMSO), and PZA (400 µg ml^-1^). The following post-infection (p.i.) time points were 0, 1, 2, 5, 8, and 10 days. Media containing appropriate treatments was replaced every 2 days. Upon p.i. completion, cells were washed thrice in PBS and lysed using 1.0 mL of 0.1% Triton-X 100 solution. Cell lysates were collected, pelleted, and washed thrice with PBS to remove detergent. Pellets were resuspended in 1.0 mL of Middlebrook (MB) 7H9 complete broth containing 0.05% Tween 80 and 10% OADC. Serial dilutions were created and plated on MB7H10 complete agar containing 10% OADC. Plates were allowed to incubate for 2-3 weeks at 37°C. Colony-forming units (CFU) were recorded and converted to CFU/mL. Each treatment condition and time point were conducted in technical duplicates with three independent biological replicates.

### CellRox Green Oxidation Assay

Intracellular ROS production was measured using CellRox Green reagent (Thermo Fisher). CellRox Green reagent is a fluorescent indicator dye, which excites upon oxidation in the presence of double-stranded DNA. We followed the CellRox green method described by Coulson et al.(Coulson, Johnson et al. 2017), which was optimized for ROS detection in *M. tuberculosis*. Briefly, log phase *M. tuberculosis* H37Rv was subcultured in MB7H9 complete (pH 6.8) containing 10% ADS with PZA (100-200 µg/mL) or equivalent volume of DMSO at an OD_600nm_ of 0.2 for 2 days at 37°C with shaking. *M. tuberculosis* cultures were subsequently treated with varying concentrations of H_2_O_2_ (0.5-10 mM) for 0.5, 2, 6 and 24 hrs at 37°C with agitation. CellRox Green reagent (5 µM) was added to each culture prior to the last 0.5 hrs of incubation. Subsequently, *M. tuberculosis* cells were pelleted, washed twice with PBS, and fixed in 10% formalin for 2 hrs at room temperature (RT). *M. tuberculosis* cultures were then washed twice as before, re-suspended in 0.3 mL of PBS, and transferred to a 96 well black microtiter plate. Fluorescence (excitation 485 nm, emission 520 nm) and growth was measured on a Synergy H1 hybrid reader (BioTek) and Gen5 2.01 software. Results were normalized to input cellular material determined by OD_600_. All treatments were conducted in duplicate. The intracellular ROS assay was performed thrice.

### Graphs and Statistical Analysis

All figures were created in GraphPad Prism ver. 5 (GraphPad Software, La Jolla, CA). Two-way analysis of variance (ANOVA) with Bonferroni correction was performed on data using the analysis tool in GraphPad Prism software. Results with a *P <* 0.05 were considered statistically significant.

## Acknowledgements

This study was supported by funds from the National Institutes of Health (AI123146 to A.D.B.). N.A.D. was supported by training grant HL007741. E.A.L. was supported by a National Academies of Sciences, Engineering and Mathematics Ford Foundation Postdoctoral Fellowship and a NIH diversity supplement award AI123146-S01. We thank the staff of the University of Minnesota BSL-3 program for providing support for these studies.

